# Predicting Protein Producibility in Filamentous Fungi

**DOI:** 10.1101/138560

**Authors:** Karmen L Dykstra, Juho Rousu, Mikko Arvas

## Abstract

In this paper we study the problem of predicting the producibility of recombinant proteins in filamentous fungi, especially *T. reesei*, using machine learning methods. We train supervised and semi-supervised support vector machines with protein sequences, represented by their amino acid composition as well as protein family and domain information. Our results indicate, somewhat surprisingly, that quite modest amount of proteins with experimental data are required to build a state-of-the-art classifier and that additional unlabeled sequences in semi-supervised models do not bring increased predictive performance. Our experiments in cross-species prediction show that models trained for the filamentous fungus *A. niger* protein dataset can be generalized to predict protein producibility in *T. reesei*, and vice versa, without sacrificing too much accuracy, regardless of their approximately 500 millions years of divergence. However, predictors trained on *E. coli* and *S. cerevisiae* datasets gave variable performance when applied to the filamentous fungi datasets, indicating that while protein producibility prediction can be generalized accross related species, fully generic prediction tools applicable to any protein production host may not be realistic to achieve.

## 1 Introduction

Industrial recombinant protein production has wide applications in life sciences, medicine and industry. Microbial hosts capable of producing high amounts of proteins are needed to make protein production economically viable. Filamentous fungi species are highly suitable as hosts for industrial recombinant protein due to their inexpensive growth requirements and innate ability to secrete proteins in large quantities. In filamentous fungus *Trichoderma reesei*, for example, the production of some endogenous proteins can be as high as 100 g/L [1]. However, recombinant protein production experiments are prone to fail, in which case no recombinant protein product is detectable in the extracellular medium. In order to increase the success rate of laboratory experiments, it is of strong interest to develop a computational tool that takes a protein sequence of interest as input, and returns a prediction of its success or failure of production in the intended host species. Such tools preemptively identify problematic sequences and recommend promising candidates for production from sequence databases.

Computational methods for protein producibility have been developed since 1991, when Wilkinson and Harrison introduced a statistical measure to estimate the expected solubility of proteins upon expression in *E. coli* [2], and several tools are currently available for recombinant protein production in *E. coli*. These tools are trained to predict *protein solubility* upon expression because this is a significant point of failure for protein experiments in *E. coli*. The prominence of *E. coli* as a production host, and the ready availability of large datasets of protein sequences with an experimentally verified solubility status, have contributed to a proliferation of such methods [3], some of which have been implemented as publicly available webtools [4]. Two recent and high-performing methods include PROSO II [5] and ccSOL *omics* [6]. PROSO II implements a two-level logistic regression classifier trained on the output from a Parzen window and k-mer features, while the ccSOL *omics* model uses Fourier coefficients and an artificial neural network classifier.

The first computational results for recombinant protein production in fungal host cells have only appeared in the last decade, to predict protein expression in the yeasts *Pichia pastoris* [7] and *Saccharomyces cerevisiae* [8], and protein secretion in the filamentous fungus *Aspergillus niger* [9, 10]. Van den Berg et al. [9] compared 9 different classifiers and found that Linear Discriminant Analysis (LDA) and Support Vector Machines (SVMs) yielded the highest performances on a dataset of homologous sequences tested in *A. niger* [9]. In a subsequent work, van den Berg et al. demonstrated a classifier performance of 85% Area Under Curve (AUC) on homologous sequences, and 75% AUC on heterologous sequences in *A. niger* [10]. They combined a linear kernel SVM with an extensive comparison of sequence-based features, including features calculated from the DNA sequence, amino acid sequence, predicted solvent accessibility, and predicted secondary structure. The authors implemented a linear kernel SVM trained on amino acid composition as a publicly available webtool, HIPSEC ^[1]^.

In this paper, we tackle the problem of predicting protein production in an important industrial protein production host, *Trichoderma reesei*, which is a filamentous fungus, like *A.niger*. Building on van den Berg’s work predicting protein producibility in *A. niger*, we train Support Vector Machines (SVMs) using amino acid composition features on datasets collected from *A. niger* and *T. reesei*. Besides the amino acid composition, we explore other representations of proteins, namely, protein family and domain information. The motivation is to study whether the proteins contain conserved protein domains whose presence is correlated with their producibility.

We additionally seek to use information from unlabeled protein homologs to improve producibility classification. The concept of using unlabeled data in combination with a limited labeled dataset is broadly termed *semi-supervised learning*, and has been successful in a range of applications including natural language processing and text classification [11]. These methods are of special interest for the case of training protein producibility predictors because labeled data is expensive and time-consuming to collect, but millions of unlabeled protein sequences are instantly available from online databases. In this work, we report experiments on a semi-supervised extension of SVMs, the Transductive SVM (TSVM) [12, 13].

## 2 Methods

### 2.1 Data

We used three different datasets of protein sequences, aligned by the host species in which they were tested for protein producibility.

- The *Aspergillus niger* dataset [9, 10] consists of 345 homologous sequences (178 positives, 167 negatives) tested for successful secretion. Each sequence was inserted into a vector with the strong glucoamylase promoter (*PGlaA*), and the modified cells were grown in shake flasks, filtered, and put on SDS-PAGE gel, where the detection of a visible band in the gel was defined as successful production [9]. To avoid sequence redundancy, BLASTCLUST [14] was used to cluster sequences with greater than 40% identity, and only one sequence from each class was kept for each cluster.
- The *Trichoderma reesei* datasets consist of both heterologous and homologous sequences and are internal to VTT Technical Research Centre of Finland Ltd. The heterologous dataset of sequences tested in the host *T. reesei* consists of 19 sequences (10 positives, 9 negatives) from a variety of species. These were tested at different times over the course of many years, with different experimental protocols. The homologous *T. reesei* dataset consists of 31 sequences (31 positives, 0 negatives) expressed under the same experimental protocol and deemed to be successfully produced according to their detection in protein gels. These sequences were selected for their high likelihood of being secreted according to previous proteomics experiments [15], and include known highly produced cellulases such as CBHI and CBHII. For model training, the heterologous and homologous sequences were combined. The heterologous, homologous, and combined *T. reesei* datasets were individually clustered using CD-HIT [16, 17] using a 60% sequence identity threshold. One sequence from each class was kept from each cluster. The CD-HIT filtered *T. reesei* dataset with both heterologous and homologous sequences contains 49 sequences (40 positives, 9 negatives).
- The *Saccharomyces cerevisiae* dataset consists of 2000 homologous sequences (1000 positive, 1000 negative) originally from [18] and BLASTCLUST [14] filtered by [8]. These sequences come from an analysis of the yeast proteome, wherein each ORF was tagged with a highaffinity epitope and expressed from the natural chromosomal location. Proteins expressed during log-phase growth were then measured through immunodetection of the common tag. After BLASTCLUST filtering, the sequences were ordered by the detected levels of protein molecules and the top and bottom 1000 were respectively given positive and negative labels. For the training of a semi-supervised prediction model one uses a combination of labeled and unlabeled data. For unlabeled sequences we used two data sources:
- The UniProt SwissProt database from October of 2013, consisting of over 500,000 unlabeled protein sequences, was used for preliminary experiments for finding good cut-off levels for sequence similarity. We used the Basic Local Alignment Search Tool (BLAST) [19] [20] to search for unlabeled sequences with varying degrees of sequence similarity to the labeled sequences. We used the *E*-value score returned by BLAST as the measure of sequence similarity. *E*-value represents a corrected *P*-value of the sequence similarity score, that is, the likelihood to find as high sequence similarity as the observed value by random. The correction counteracts the multiple testing of a query sequence against a large sequence database. We performed a BLAST of the *A. niger* dataset against the UniProt SwissProt sequences, and selected all sequences that were within a BLAST *E*-value of 10 to the labeled dataset. This returned 19,845 sequences. A histogram of these sequences by BLAST *E*-value to the closest labeled sequence is shown in Figure 1. There are a greater number of proteins with weaker homology to the labeled sequences, as shown by the increase in number of sequences as the *E*-value increases.
- The UniProt TrEMBL database, which contains over 54 million sequences, was used as the data source for the main experiment with semi-supervised models, queried with sequence similarity cut-off levels established with the SwissProt dataset.

**Figure 1.**
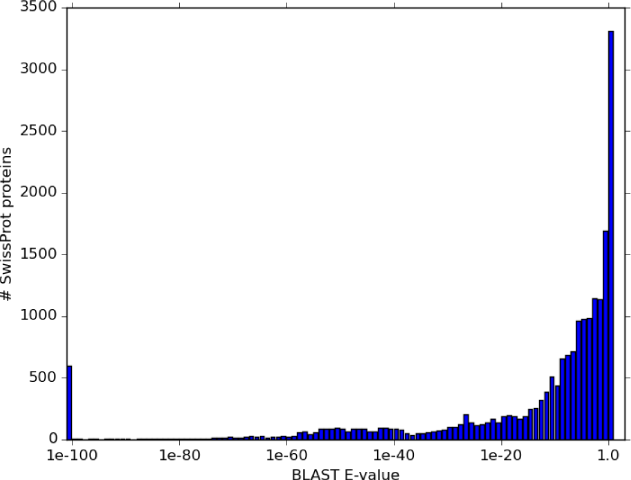
Histogram of number of sequences vs. smallest BLAST *E*-value to the labeled *A. niger* sequences.

### 2.2 Feature representations

We represented each protein sequence by its normalized amino acid composition. Define *ni* as the number of times that amino acid *i* occurs in the input protein sequence. Using the 20 standard amino acids, each feature vector can be defined as **x** ϵ ℝ^20^ = (*x*_1_, …, *x*_20_) where the features 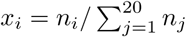 denote the relative frequency of the amino acids.

In addition, we used the InterProScan tool ^[2]^ [21] to build a feature representation based on protein-specific signatures, to test for the efficacy of protein family and domain information in predicting protein production in *A. niger* and *T. reesei*. We used all InterProScan signatures that were detected in at least two labeled proteins in the union of the *A. niger* and *T. reesei* combined datasets, which amounted to 391 signatures. A binary feature vector **x** ϵ {0, 1} = {*x*_1_, …, *x*_391_} was constructed for each protein sequence by placing a 1 if the signature was detected in the protein, and a 0 otherwise.

### 2.3 Support Vector Machines

We applied support vector machines (SVMs) to classify protein sequences because they have been shown to perform favorably for predicting producibility in filamentous fungi when compared to other methods [9]. In our experiments, the SVMs were trained with the Scikit-Learn [22] package.

The SVM is a widely used pattern recognition model that learns a linear decision surface through a feature representation of the input data points. The SVM objective is to minimize a combination of the training error, which describes the model’s fit of the data, and a regularization term, to avoid overfitting. The objective can be alternatively interpreted as maximizing the minimum distance, called the margin, between examples of either class, whilst allowing some outliers (examples too close to the other class).

The formal definition of the SVM from [23] is as follows. We are given a set of *L* training pairs *𝓛* = {(**x**_1_, *y*_1_), …, (**x**_*L*_, *y*_*L*_)}, **x** ϵ ℝ^*n*^, *y* ϵ {−1, 1}. An SVM learns a decision function

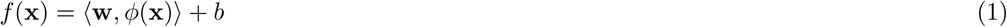

where *φ* (*·*) is the chosen feature map and 〈*·*, 〉 denotes the inner product. To return a binary prediction, the value of the function (1) is thresholded at zero. The weights **w** and the intercept *b* are learned by solving

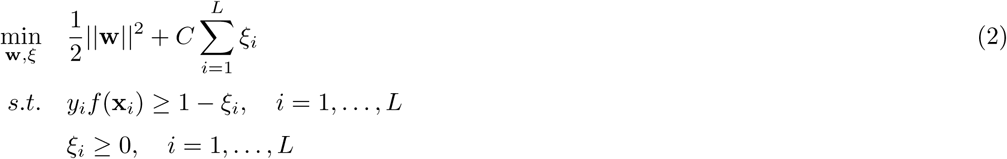

In the above equations, *ξ*_*i*_ is the hinge loss function *H* on sample *i*, defined as

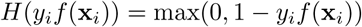

The parameter *C* in the above optimization problem represents a trade-off between the margin, controlled by ||**w**||, and the size of the error terms *ξ*_*i*_. Optimization is carried out over the quadratic programming problem formed from the dual form of the function, resulting from the Lagrangian [23]. Introducing the dual variables {*α*_*i*_, …, *α*_*L*_}, the learned decision function can equivalently be written

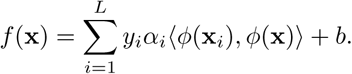

In fact, the inner product of the feature vectors 〈*φ* (**x_*i*_**), *φ* (**x**)〉 is the only input needed to learn the decision function. This inner product can be calculated directly via the *kernel function*. The choice of kernel defines the shape of the learned decision function in the input space. In this work we experimented with the following kernels:

- The linear kernel, defined by the inner product of the original real-valued vectors:

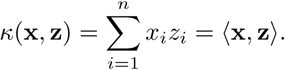 The feature map implicitly used for calculating the linear kernel is the same as the original vector, or equivalently, *φ* (**x**) = **x**.
- The RBF kernel, also known as the Gaussian kernel, is defined as

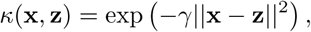

where the parameter *γ >* 0 controls to the width of the Gaussian distribution: large values correspond with “peaky” Gaussians, while small values of *γ* produce wider Gaussian kernels.
- A polynomial kernel with degree *k* is defined as

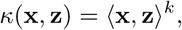

where the integer parameter *k ≥* 1 controls the shape of the decision boundary: large values of *k* allow more complicated decision boundaries than small values.

### 2.4 Semi-supervised SVM models

We investigated the application of a semi-supervised SVM model, the transductive SVM (TSVM) [13], to the problem of predicting successful protein production in filamentous fungi. The TSVM extends the SVM framework to make use of unlabeled examples to improve generalization accuracy based on the premise that there are “gaps” between the data points belonging to different labeled classes. This echoes the reasoning behind the margin maximization performed by the original SVM model, but the TSVM extends the idea by using unlabeled data points to locate where to place the decision boundary so as to maximize the distance between classes. In practice, this principle takes the form of an additional term in the optimization function that penalizes decision functions that place unlabeled data points in the margin, close to the decision boundary.

With the addition of *U* unlabeled examples *U* = {*x*_*L*+1_, …, *x*_*L+U*_}, and keeping the same notation as used for the fully-supervised SVM (Equation 2), the objective function can be formulated as

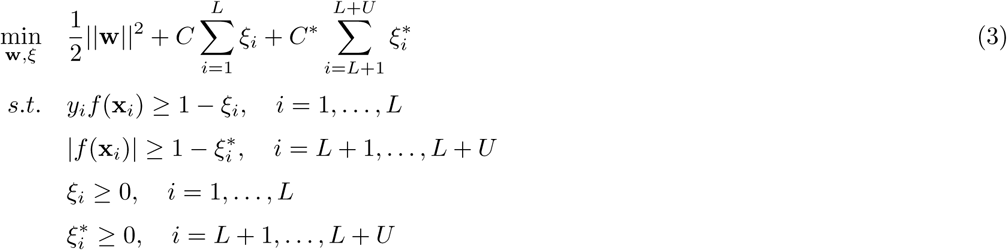

We used the TSVM implementation by Collobert et al. [13] ^[3]^.

### 2.5 Evaluation metrics

To evaluate the predictive performance of models, we used two evaluation metrics:

- Balanced accuracy, calculated by taking the average of the True Positive Rate (TPR, the fraction of positive examples correctly predicted to be positive) and True Negative Rate (TNR, the fraction of negative examples correctly predicted to be negative): 0.5 *·* (*T P R* + *T N R*). The equal weighting of accuracy within each class is useful for evaluating on data with an unbalanced number of examples from each class, as is the case with the combined *T. reesei* dataset.
- Area Under ROC Curve (AUC), which measures the accuracy of the model *f* (**x**) *≥ σ* to classify the data when the threshold *σ* of the decision function is varied. Alternatively, AUC can be seen as measuring the model’s ability to correctly order or rank the sequences. An AUC of around 0.5 means the classifier ranks the output no better than a random ordering and 1.0 means the classifier perfectly ranks the output sequences so that they can be separated by true class.

## 3 Results

### 3.1 Comparison with Existing Tools

In order to assess the generalization performance of predictors across different species of industrial protein producers, we tested the performance of different models, some of which are available as webtools, to predict class labels for the *A. niger, T. reesei*, and *S. cerevisiae* datasets, for which the resulting accuracies are shown in Table 1. Predictions from PROSO II, ccSOL *omics*, and HIPSEC were gathered directly from the publicly available websites for these tools. For PROSO II, positive and negative labels were taken directly from the output, and for ccSOL *omics* and HIPSEC the returned predictions were respectively thresholded at 50 and 0. The *T. reesei SVM* and *S. cerevisiae SVM* models refer to linear kernel SVMs that we trained on the respective datasets, using the amino acid composition of the sequences as training features. SVM parameters were set by a 5-fold crossvalidation loop for the *T. reesei SVM* model, and a 10-fold cross-validation loop for the *S. cerevisiae SVM* model. Such a model was previously shown to yield a 0.79 average cross-validation AUC on the *S. cerevisiae* dataset [8].

**Table 1.** Accuracies of 5 models for predicting class labels for *A. niger, T. reesei* homologous (hom) and heterologous (het) components, and *S. cerevisiae* datasets. The results for PROSO II [5], ccSOL *omics* [6] and HIPSEC [10] were gathered from the tool websites. Performance of the model on the training data is omitted (-).

| Tool | *A. niger* [10] | *T. reesei* hom | *T. reesei* het | *S. cerevisiae* |
| --- | --- | --- | --- | --- |
| PROSO II | 0.52 | 0.68 | 0.53 | 0.58 |
| ccSOL *omics* | 0.50 | 0.55 | 0.63 | 0.48 |
| HIPSEC | - | 0.81 | 0.84 | 0.50 |
| *T. reesei* SVM | 0.70 | - | - | 0.53 |
| *S. cerevisiae* SVM | 0.57 | 0.94 | 0.63 | - |

### 3.2 Training an SVM with Few Labeled Training Examples

The HIPSEC model, consisting of a linear kernel SVM trained with the *A. niger* dataset, performed well on the *T. reesei* data, returning balanced accuracies of 0.84 and 0.81 on the heterologous and homologous components, respectively. In this case, the training dataset consists of 345 sequences. Given that it is costly to gather this amount of experimental data, we investigated whether a similar performance could be reached with fewer training examples.

We first assigned the data to 10 random cross-validation folds. For each fold, we held the test examples constant for all iterations, but decreased the number of training examples by 10 in every iterations were continued until 10 training examples remained from each fold. We used stratified sampling so that the ratio of positive to negative class training examples is the same in each subsample.

Models were trained with RBF and linear kernels. *C* and *γ* parameters were selected separately for each fold using a second 5-fold cross-validation loop within the training data, such that the parameters maximized the average accuracy across the 5 inner folds. Parameter selection was performed once in the first iteration using all labeled training data, and these parameters were used for all the following iterations. The average performance across folds and its standard deviation are shown for each iteration in Figure 2. Repetition of the experiment with different randomized data splits yielded results similar to those shown.

**Figure 2.**
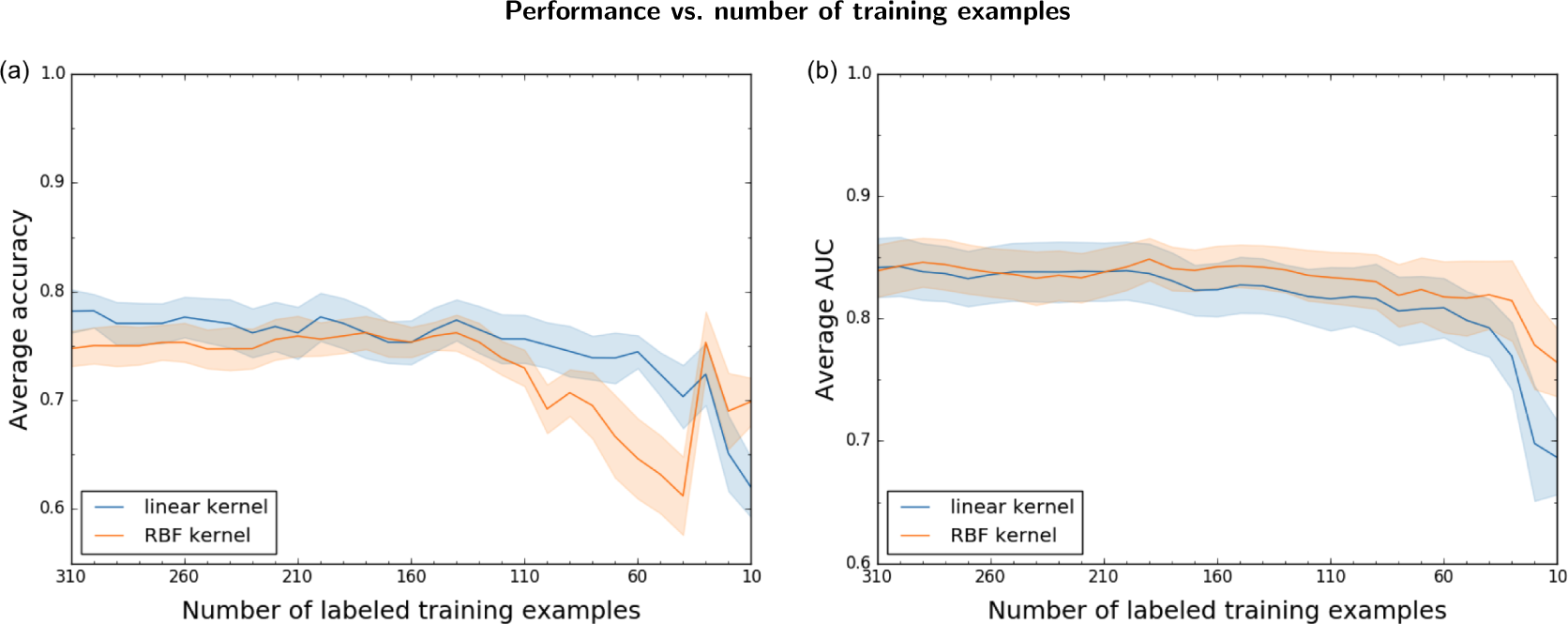
Accuracy (pane a) and AUC (pane b) as a function of the number of labeled training examples for an SVM trained with the *A. niger* data. Solid lines show the average across 10 cross-validation folds and the shaded area is the standard error. The blue and orange colors show results using a linear and RBF kernel, respectively.

### 3.3 SVM performance on different feature sets and kernels

We compared the performance of different features and model choices for training and testing on the *A. niger* and *T. reesei* datasets. In addition to training SVM models with normalized amino acid composition as input features, we tested the contribution of protein characteristics to the prediction problem by incorporating InterProScan features (Section 2.2), either by concatenating the feature vectors prior to training the models, or by taking the element-wise kernel product of the normalized kernel matrices calculated from each feature set. We tested linear, RBF, and polynomial kernels for every feature combination.

We trained and tested SVM models on both the *A. niger* and combined *T. reesei* datasets. We used 10-fold cross-validation to train and evaluate models trained with the *A. niger* dataset, and leave-one-out cross-validation to evaluate the *T. reesei* model, because of its smaller size. For the *A. niger* model, we calculated the average AUC and balanced accuracy across cross-validation folds. For the *T. reesei* model, we calculated the balanced accuracy of the aggregated leave-one-out predictions.

AUC was not calculated for the *T. reesei* models because the distances of individual samples from the classification hyperplane, returned by leave-one-out cross-validation, are not directly comparable across models due to different selected regularization parameters. In addition to testing model performance within each dataset by cross-validation, we tested the generalization performance across datasets. Features and kernel choices that generalize well may capture protein characteristics that indicate protein producibility in filamentous fungi at a broader scale than within individual species. The SVM *C* parameter and kernel parameters were set using a nested cross-validation scheme, with a 3-fold inner cross-validation loop over the training data for each fold. The parameters that resulted in the highest average balanced accuracy across the inner cross-validation loops were selected. In these experiments the *C* parameter and *γ* parameter for the RBF kernel were set using a grid search over {10^−3^, 10^−2^, 10^−1^, 10^0^, 10^1^, 10^2^, 10^3^} and the *k* parameter for the polynomial kernel was set with a grid search over {2, 3, 4, 5, 6}.

Results for the SVM models trained on the *A. niger* and *T. reesei* datasets are shown in Table 2 and Table 3, respectively. The type of kernel is shown in parentheses, and the highest value achieved in each column is in bold font. For the *A. niger* dataset, where the average was taken across crossvalidation folds, its standard deviation is shown in parentheses. The first observation is that aminoacid composition is clearly the dominating feature type, and that non-linear kernels do not increase accuracy. The kernel product seems to be overall the best-performing method of integrating the Interpro features with the amino acid composition; however, even that combination fails to perform at the level of amino acid features alone. The non-linear RBF and polynomial kernel do not have much effect on the *A. niger* trained model. On the model trained with *T. reesei* data, the non-linear kernels actually hurt the performance.

**Table 2.** Performance of feature/kernel pairs for SVM model trained with **A. niger** data. *A. niger* and *T. reesei* column headers indicate the dataset used for evaluation. The *A. niger* column shows the cross-validation average with its standard deviation in parentheses.

|  | *A. niger* |  | *T. reesei* |  |
| --- | --- | --- | --- | --- |
|  | AUC | Bal Acc | AUC | Bal Acc |
| AA composition (linear) | **0.84 (0.08)** | 0.76 (0.09) | 0.86 | 0.84 |
| InterProScan (linear) | 0.74 (0.12) | 0.67 (0.12) | 0.83 | 0.68 |
| Feature concatenation (linear) | 0.75 (0.10) | 0.68 (0.08) | 0.78 | 0.60 |
| Kernel product (linear) | 0.78 (0.11) | 0.70 (0.08) | 0.81 | 0.76 |
| AA composition (RBF) | **0.84 (0.07)** | **0.77 (0.08)** | 0.83 | 0.73 |
| InterProScan (RBF) | 0.75 (0.12) | 0.67 (0.12) | 0.74 | 0.69 |
| Feature concatenation (RBF) | 0.75 (0.11) | 0.67 (0.11) | 0.80 | 0.61 |
| Kernel product (RBF) | 0.76 (0.10) | 0.68 (0.11) | 0.76 | 0.62 |
| AA composition (polynomial) | 0.82 (0.09) | **0.77 (0.09)** | **0.89** | **0.88** |
| InterProScan (polynomial) | 0.75 (0.11) | 0.68 (0.12) | 0.78 | 0.60 |
| Feature concatenation (polynomial) | 0.74 (0.11) | 0.68 (0.11) | 0.79 | 0.60 |
| Kernel product (polynomial) | 0.77 (0.11) | 0.70 (0.10) | 0.82 | 0.69 |

**Table 3.** Performance of feature/kernel pairs for SVM model trained with **T. reesei** combined data. *A. niger* and *T. reesei* column headers indicate the dataset used for evaluation. The *T. reesei* column shows the balanced accuracy for leave-one-out class predictions.

|  | *A. niger* |  | *T. reesei* |
| --- | --- | --- | --- |
|  | AUC | Bal Acc | Bal Acc |
| AA composition (linear) | **0.75** | 0.69 | **0.80** |
| InterProScan (linear) | 0.67 | 0.51 | 0.60 |
| Feature concatenation (linear) | 0.73 | 0.51 | 0.66 |
| Kernel product (linear) | 0.72 | 0.51 | 0.63 |
| AA composition (RBF) | 0.74 | 0.67 | 0.62 |
| InterProScan (RBF) | 0.51 | 0.51 | 0.70 |
| Feature concatenation (RBF) | 0.51 | 0.51 | 0.63 |
| Kernel product (RBF) | 0.51 | 0.51 | 0.67 |
| AA composition (polynomial) | 0.74 | **0.70** | 0.65 |
| InterProScan (polynomial) | 0.69 | 0.51 | 0.59 |
| Feature concatenation (polynomial) | 0.72 | 0.50 | 0.60 |
| Kernel product (polynomial) | 0.72 | 0.52 | 0.54 |

### 3.4 Semi-supervised models

Semi-supervised methods use a combination of labeled and unlabeled data. Since modern sequencing technologies have made a wealth of protein sequences publicly available, the question immediately arises of *how to select* the protein sequences that form the unlabeled dataset.

We conducted preliminary experiments to determine the effect of sequence distance in BLAST *E*-value space on performance using the semi-supervised TSVM model. We separated the 19,845 unlabeled SwissProt sequences into 5 bins using the following *E*-value thresholds: {10^−60^, 10^−20^, 10^−1^, 10^0^}. We performed 5 rounds of 10-fold cross-validation with a linear kernel TSVM trained on amino acid composition features. In each round we used the same labeled *A. niger* examples for training, but different unlabeled sequences. In round *k* = 1, …, 5, we sample 1000 unlabeled sequences uniformly at random from the first *k* bins, where the bins are ordered by increasing *E*-value thresholds. Thus the number of unlabeled sequences is kept constant, but the maximum BLAST *E*-value is increased in each round. The resulting AUC values are shown in Table 4. Based on Table 4, the highest performances of the TSVM model resulted when using sequences from the first two bins. We thus limited unlabeled sequences to those with BLAST *E*-value smaller or equal to 10^−20^.

**Table 4.** AUC values obtained by 10-fold cross-validation of a linear TSVM trained on *A. niger* dataset using amino acid composition features, using 1000 unlabeled sequences. Each AUC values corresponds to a given maximum BLAST *E*-value threshold against labeled *A. niger* sequences.

| max *E*-value | $10^{-60}$ | $10^{-20}$ | $10^{-1}$ | $10^0$ | $10^1$ |
| --- | --- | --- | --- | --- | --- |
| AUC | 0.829 | 0.817 | 0.794 | 0.792 | 0.786 |

For subsequent experiments, we switched from the SwissProt database to the larger UniProt TrEMBL database. In order to keep sequence similarity threshold consistent, we used the SwissProt database size *N* = 542, 901 as the normalizing factor in the *E*-value, rather than the TrEMBL database size. We note that this rescaling of *E*-values does not constitute a statistical problem, since we do not use the *E*-values to make claims of statistical significance. BLAST of the labeled *A. niger* and *T. reesei* sequences against the TrEMBL database returned 307,301 unlabeled candidate TrEMBL sequences for the *A. niger* dataset and 72,679 sequences for the *T. reesei* dataset.

We evaluated the performance of a linear kernel TSVM (Section 2.4) and two different sampling procedures to select the unlabeled sequences to use during training.

- In the “random” sampling method, we selected *U* ∈ {500, 2000} unlabeled sequences for each cross-validation fold from the TrEMBL sequences with *E*-value at most 10^−20^ and combined them with the labeled training data
- The “close” method, sampled from TrEMBL sequences *k* nearest neighbors of each labeled measure. We used *U* ∈ {500, 2000} as the number of unlabeled sequences. The number of nearest-neighbors to select for each labeled sequence was set equal to *k* = ┌*U*/*N*┐, where *N* denotes the number of labeled examples and ┌·┐ rounds up to next larger integer.

For labeled training examples, we examined two selection strategies: training with all labeled training examples, and training with a subset of 10 labeled training examples. These 10 examples were randomly selected from each fold, such that the ratio of negative and positive examples in the subsample was the same as in the complete dataset.

The *C* parameter for the SVM was set once using all labeled examples. The best parameters for each cross-validation fold were selected as the parameters that maximized the average balanced accuracy within an inner 3-fold cross-validation loop. These parameter values, set once for each dataset, were applied for all semi-supervised experiments, including those with only 10 labeled training examples.

For the *A. niger* dataset, the regularization parameter for the TSVM, *C*^*^, was selected by an inner 3-fold cross-validation loop, so that the selected value of *C*^*^ maximized average accuracy across folds. A grid search over 10^*i*^ for *i* ∈ {−3, −2, −1, 0, 1, 2, 3} was performed to select *C*^*^. The parameter was kept at its default value for the *T. reesei* dataset, because of the dataset’s small size.

The results of this experiment are shown in Table 5. When all labeled training examples are used, the unlabeled examples do not bring any advantage. On *A. niger* the balanced accuracy is unaffected by the unlabeled data, while for *T. reesei*, the balanced accuracy goes down upon adding unlabeled examples. On the other hand, in the case when only 10 labeled examples are used, TSVM is able to improve balanced accuracy. The “close” selection strategy seemed to be better than “random” on *T. reesei* data, however on *A. niger* data the different strategies performed equally.

**Table 5.** Balanced accuracy performance of a TSVM with a linear kernel. Shows avg. (std.) for *A. niger* model and leave-one-out balanced accuracy for *T. reesei* model.

|  | *A. niger* |  | *T. reesei* |  |
| --- | --- | --- | --- | --- |
|  | All ex | 10 ex | All ex | 10 ex |
| SVM | 0.76 (0.09) | 0.61 (0.11) | 0.80 | 0.57 |
| random 500 | 0.75 (0.07) | 0.66 (0.11) | 0.70 | 0.60 |
| random 2000 | 0.76 (0.07) | 0.67 (0.09) | 0.70 | 0.60 |
| close 500 | 0.76 (0.07) | 0.68 (0.12) | 0.77 | 0.62 |
| close 2000 | 0.76 (0.06) | 0.68 (0.10) | 0.77 | 0.62 |

### 3.5 SVM Predictions on SwissProt

The model evaluations depicted before give us estimates for how well we can predict the success of available labeled data, primarily composed of sequences from the native *T. reesei* and *A. niger* genomes. In this section we sought to understand what sequences are predicted to be “producible” at a wider scale, by analyzing SVM model predictions on the Uniprot SwissProt database of protein sequences. Here we used the SVM model trained with the linear kernel on amino acid composition. We tested for enrichment of “successful” predictions in taxonomic classification of the organism from which the sequence originates, and in enzyme function as specified by Enzyme Commission (EC) numbers.

SwissProt includes taxonomic classification of the organism from which each protein sequence originates. An example entry from the database for a plant protein sequence is as follows: Eukaryota; Viridiplantae; Streptophyta; Embryophyta; Tracheophyta,Spermatophyta; Magnoliophyta; eudicotyledons; Gunneridae, Pentapetalae; rosids; fabids; Fabales; Fabaceae; Papilionoideae,Fabeae. To separate sequences by the classification of their origin species, we defined taxonomic categories by the fifth level of their taxonomic classification, where each level is separated by semi-colons. In the example sequence taxonomy, the sequence would belong to the “Tracheophyta” category.

47% of the sequences in SwissProt also have an Enzyme Commission (EC) number. The EC number system, assigned by the Nomenclature Committee of the International Union of Biochemistry and Molecular Biology (NC-IUBMB)^[4]^ describes the reactions catalyzed by the protein. In tests for enrichment by EC class, we separated sequences by the third number in their EC number. As an example, cellulases have an EC number of 3.2.1.4, so in our enrichment tests they would be counted in the “3.2.1” category.

To test for a significantly high number of predicted “successes” per category, we used the hypergeometric statistical test, implemented as phyper in the R programming language. Given a sample population, this tests whether a sub-sample of items, taken without replacement, contains a largerthan-expected proportion of items of a particular type. In our case, the “sample population” is the SwissProt database, the sub-sample taken consists of all sequences with a positive prediction, and the “type” tested for significant presence in the sub-sample is either the taxonomy of the organism of origin or the sequence EC number.

We only considered EC number or taxonomic categories that contained at least 20 sequences in SwissProt, to avoid deeming a category as “enriched” with positive predictions if there were very few examples from which to draw this conclusion.

The results of the enrichment tests for the *A. niger* model are represented in Figure 3. The *T. reesei* model gave similar results (data not shown). Various fungal hydrolytic enzymes active on carbohydrates and peptides show the most prominent enrichments. However, enzymes of similar classes, in particular, 3.2.1-class glycosylases, i.e. cellulases, are enriched in non-fungal eukaryotic groups also.

**Figure 3.**
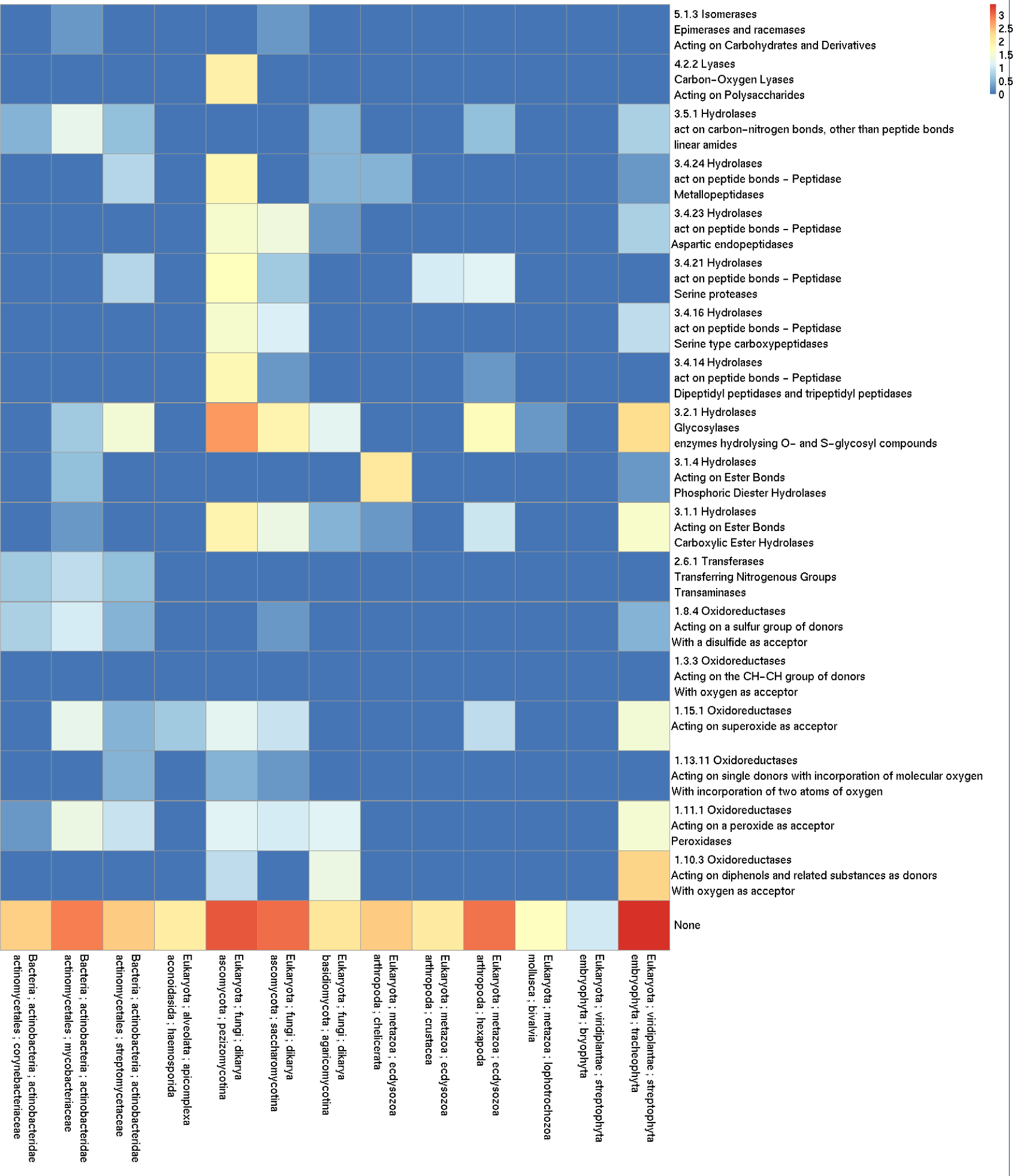
EC number/ organism taxonomy enrichment for the linear kernel model trained with the *A. niger* dataset. Cell coloring is a function of the base 10 logarithm of the number of predicted successes with the donor organism taxonomy of the cell column and EC number of the cell row. The EC numbers and organism taxonomies that form the rows and columns were enriched in the model positive predictions with a p-value of 0.

### 4 Discussion

We successfully applied an SVM trained with amino acid composition features to predict the successful production of proteins in A. niger and T. reesei. While this approach was previously applied to A. niger [55], we demonstrated its high success rate for proteins produced in a different filamentous fungus host species, T. reesei.

While the *A. niger* dataset consists of over 300 protein sequences, we demonstrate that as few as 50 sequences can be used to train a predictive model with comparable performance. This finding is further supported by the high predictive abilities of the *T. reesei* model, trained on almost 50 sequences. We applied the SVM predictors from both *A. niger* and *T. reesei* datasets to the protein sequences in UniProt SwissProt, and observe that enzyme classes with a significant portion of predicted successes include cellulases and proteases which would be expected to be produced successfully. In addition, we showed that the training data does not need to be balanced with respect to positive and negative labels, as indicated by the success of the model trained with the unbalanced T. reesei data and applied to the A. niger data.

Experiments with the semi-supervised TSVM gave mixed results. It seems that when very few labeled examples (10 were used in our example) are available, adding unlabeled examples can help somewhat. However, the advantage seemed to disappear and turn to disadvantage quite rapidly, which limits the applicability of semi-supervised models in protein producibility prediction.

In order to find alternative, hopefully more predictive representations for protein sequences over the amino acid composition features, we explored the use of family domain information to improve the model through the inclusion of InterProScan features, consisting of protein family and domain information. The rationale for testing them is that, because enzymes which are secreted naturally by filamentous fungi degrade biopolymers in the surrounding environment for use as energy sources, it is likely that secreted proteins contain conserved protein domains related to this function. The addition of protein domain information has also been found to improve performance for predicting protein subcellular location [24]. However, our experiments indicated that these features do not benefit the protein producibility task, compared to using amino acid content information only, and can actually hurt the performance. Hence, protein family and domain information should probably be excluded in future modeling efforts.

The solubility of a protein inside a cell or secretability of a protein out of cell could depend mostly on its physical properties. Also the solubility and secretability might essentially be very similar physical properties of a protein. The ability to fold efficiently and to stay solute inside a cell is at least a prerequisite for efficient secretion. Hence, prediction of solubility with mix species data sets has been proposed [25]. Nevertheless the intracellular conditions at least between Fungi and Bacteria appear so different that proteins that are insoluble upon expression in *E. coli*, can be highly soluble when expressed in *Saccharomyces cerevisiae* [26].

In order to reliably interpret prediction results across different species the species specific data sets used for prediction should be somehow comparable. The data from species we use have been produced with various laboratory techniques. Most importantly the labelling of proteins to successful and unsuccessful has been done by independent qualitative choice of original authors for each data set. Hence, it is quite remarkable that a model trained on *A. niger* data works so well to predict secretion in *T. reesei*, regardless of the approximately 500 millions years of divergence between the species [27]. In contrast, a model trained on *S. cerevisiae*, an even more distant fungus, gives mixed results (Table 1). This could either point to problems in data or suggests the possibility of yet undiscovered fundamental differences between the secretion systems of Pezizomycotina (*A. niger* and *T. reesei*) and Saccharomycotina (*S. cerevisiae*) fungi.

The naturally secreted proteins of Pezizomycotina and Saccharomycotina both contain a wealth of glycosidic and peptide bond active enzymes that the fungi use to transfrom biomass into a form that is easier to internalise. However, Peziziomycotina are generally able to degrade complex polysaccharides, e.g. cellulose, with various cellulases and accessory enzymes, while Saccharomycotina only use simple mono or dimeric glycosides. These differences are clearly visible from their genomic content of enzyme genes [28].

The *A. niger* and *T. reesei* data sets used to build the models are rich in proteases and cellulases hence it is not surprising that such enzymes are predicted to be secreted in the SwissProt data. However, not only such fungal enzymes but also proteins from very distant taxons are predicted to be secreted. This could imply that specific protein families have properties that make them physically stable and hence secretable, such as tight globular structures or large numbers of sulphur bonds as seen in cellulases. In this respect it is surprising that the protein domain data performs consistently so poorly in comparison to amino acid content data.

### 5 Conclusions

In this paper we have evaluated machine learning methods in prediction of recombinant protein producibility in the filamentous fungi *T.reesei*. Our results indicate that already modest amounts of training data allow accurate predictions using a support vector machine and additional unlabeled training data did not improve the results if all labeled data was used. As input features, it was shown to be sufficient to use the amino acid content, as opposed of more elaborate protein features. Cross-species prediction was shown to be feasible between two filamentous fungi, *A. niger* and *T-reesei*, but not between more distant microbes.

We note, however, The current results, where a relatively simple SVM classification model perform better than more elaborate alternatives, may reflect a ’local optimum’ in protein producibility prediction—perhaps significantly larger datasets, that map more densely the space of producible recombinant proteins for sets of host species, would allow us to leverage more elaborate features and machine learning schemes, and let us dissect the differences between secretion systems in different hosts. This could significantly improve the production of heterologous proteins in established production systems.

## Author’s contributions

KLD and JR conceived the computational approaches for the paper. MA provided biological expertise and data. KLD performed data analysis in collaboration with JR and MA. KLD, JR and MA wrote the manuscript.

## Acknowledgements

This work has been supported in part by the European Union FP7 Cooperation Work programme (Grant BIOLEDGE FP7-KBBE-289126 ‘BIO knowLEDGe Extractor and Modeller for Protein Production’) and the Finnish Funding Agency for Innovation TEKES under the Large Strategic Opening project ’Living Factories’ (decision number 40128/14).

## Footnotes

[1] http://helix.ewi.tudelft.nl/hipsec/home.py

[2] http://www.ebi.ac.uk/Tools/pfa/iprscan5/

[3] http://mloss.org/software/view/19/

[4] http://www.chem.qmul.ac.uk/iubmb/enzyme/

